# Persistence: Using Protein Turnover to Expand the Applications of Transcriptomics

**DOI:** 10.1101/2020.07.02.185462

**Authors:** Marissa A. Smail, James K. Reigle, Robert E. McCullumsmith

**Author notes:** Corresponding author: Marissa A. Smail, University of Cincinnati, Department of Pharmacology and Systems Physiology, 2170 E. Galbraith Rd. Bldg E. Room 216, Cincinnati, OH 45237-0506, United States. Co-first authors (Marissa A. Smail, James K. Reigle).

## Abstract

One of the major issues with RNA sequencing is the lack of reproducibility between RNA and protein expression. Transcriptomics offers a holistic view of the molecular landscape of a tissue at an RNA level. However, RNA and protein expression are often at odds when measured in the same sample, raising the question whether or not changes in RNA expression translate to functional differences. This problem creates a need to devise a way to approximate protein abundance from transcriptomics data, in order to create a more complete picture of the functional landscape of a tissue. One additional measure that could be useful here is protein turnover or half-life. Once RNA is transcribed into protein, that protein can either be quickly degraded or remain in the cell for an extended period of time. The longer a protein’s half-life, the more influence it can have on its surroundings. Recently, a study used stable isotope labeling in mammals (SILAM) in combination with mass spectrometry to determine the turnover ratio of ∼2200 protein in mouse synaptosomes. This data offers a valuable opportunity to integrate protein turnover with RNA expression to gain deeper insight into the functional meaning of RNA expression changes. Here, we present the concept of this combination of protein turnover and RNA expression, which we coined as persistence. We then demonstrate the application of persistence using schizophrenia (SCZ) transcriptomics datasets. Calculating persistence for these datasets greatly improved our ability to predict protein expression from RNA expression. Furthermore, this approach successfully identified persistent genes and pathways known to have impactful changes in SCZ. These results suggest that persistence is a valuable metric for improving the functional insight that can be gained from transcriptomics data.

## INTRODUCTION

The use of RNA sequencing (RNAseq) to assess global changes in gene expression has grown rapidly in the last decade [1–3]. RNAseq allows researchers to collect transcriptomic signatures in their tissue and disease of study, yielding extensively more data than more traditional methods such as qPCR. This comprehensive view of the molecular landscape has been pivotal to uncovering novel disease mechanisms and targets. As the cost of RNAseq drops and the technology improves, it will be a central tool in biological studies moving forward [3–5].

While RNAseq is a powerful tool, it has one major caveat, in that RNA expression does not necessarily match protein expression. RNA may be present in a cell, but we have no way of knowing the rate at which that RNA is translated into protein or if that protein is active in the cell. This disequilibrium between RNA and protein suggests that RNAseq results could be misleading and fail to reveal functional insights into cell function and disease mechanisms [5–8]. While proteomics is an option, RNAseq is more commonly used and cost-effective, so finding a way to extend RNAseq applications to gain insight into protein expression would allow use of existing mRNA datasets to gain a deeper understanding of diseases or model systems [8].

Here, we propose a novel method of using protein turnover ratios to infer protein expression from RNAseq and microarray data. Every protein has a particular half-life, and therefore exists in a cell to exert its functions for a relatively predictable amount of time [9,10]. We propose that using this information about protein half-life in tandem with transcriptomics data could offer insights into protein expression. Proteins that are degraded quickly may show low protein expression even if there is a large amount of RNA present, and proteins that last may show high protein expression even if there is lower RNA. We coined the term “persistence” to define this relationship between RNA expression, protein half-life, and protein expression, and propose that analyzing persistence could offer greater functional insights into traditional RNAseq datasets [11,12].

We utilized existing protein turnover data in combination with schizophrenia (SCZ) RNAseq and RNA microarrays to assess this concept and demonstrate its potential application in disease states. In a 2018 paper, stable isotope labeling in mammals (SILAM) to assess protein turnover in mouse synaptosomes and turnover ratios were determined for ∼2,200 proteins in the brain [13]. We used these data to establish persistence scores and applied these values to existing RNA datasets. Three existing RNA datasets from our lab containing control (CTL) and SCZ subjects were examined. Here we assessed (1) the relationship between RNA abundance and protein turnover ratios in CTL and SCZ; (2) whether or not there is a shift in persistence in SCZ; (3) the biological pathways associated with high and low persistence genes; and (4) the relationship between persistence and protein expression.

Our results suggest that, while there is not a direct relationship between turnover and RNA expression or a shift in persistence in SCZ, genes identified as high and low persistence have previously been implicated in the functional deficits of SCZ. Additionally, our current method improves the ability of RNA data to predict protein data, and further studies with larger datasets and machine learning could be used to further improve the persistence calculation. If we can more closely identify the relationship between RNA, protein, and turnover, this method could expand the applications of transcriptomics by allowing it to offer insight into protein function.

## RESULTS

### There is not a relationship between protein turnover ratios and RNA abundance

We started by simply looking for a relationship between protein turnover and RNA abundance with the idea that perhaps proteins that turnover faster are transcribed faster or vice versa. Regression analysis revealed that there is no relationship between turnover and RNA in CTL or SCZ in any of our datasets (Deep CTL: Adj R^2^=0.000135, Deep SCZ: Adj R^2^=0.000267, DISC1 CTL: Adj R^2^=0.000843, DISC1 SCZ: Adj R^2^=0.000672, Super CTL: Adj R^2^=0.000319, Super SCZ: Adj R^2^=-0.000231) (Fig 1). This suggests that there is a more complex relationship between RNA and protein across the transcriptome. We cannot simply assume that high turnover proteins are transcribed more, so a more complex calculation is needed to integrate these measurements in a way that is informative.

**Figure 1:**
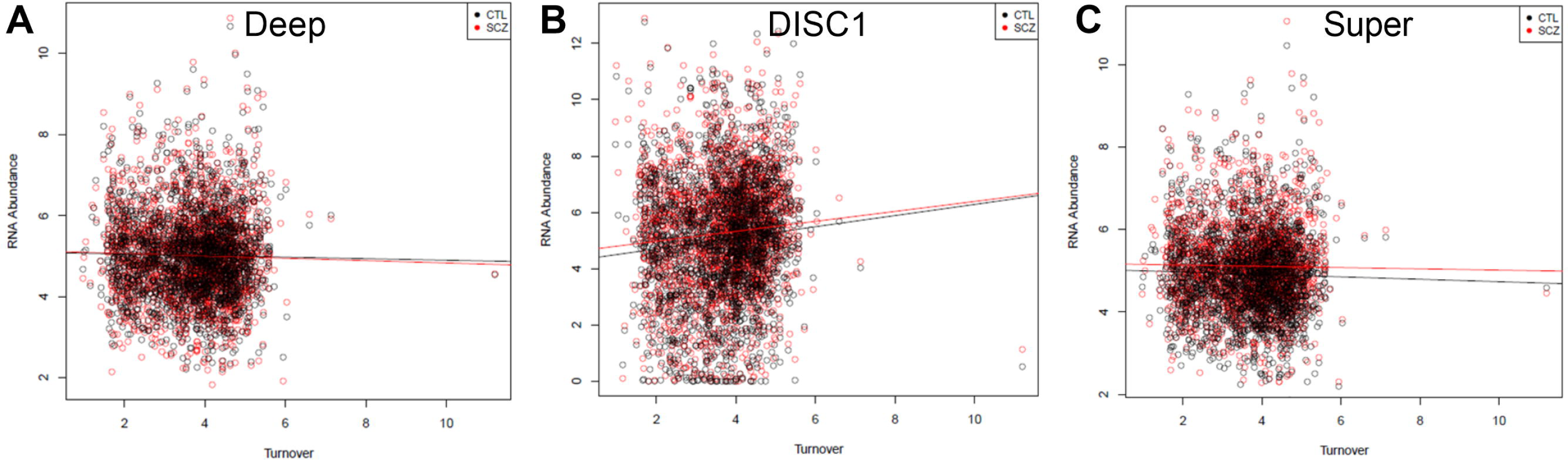
Lack of relationship between protein turnover ratio and RNA abundance. Correlation plots between protein turnover ratios and RNA abundance in (A) Deep, (B) DISC1, and (C) Super datasets. No relationships were identified in CTL (black dots) or SCZ (red dots) subjects.

### SCZ does not shift RNA expression in high or low turnover proteins

While no overall relationship was observed between turnover and RNA, it is possible that SCZ could enhance the enrichment of a certain family of turnover proteins. For example, short-lived proteins could be expressed more in SCZ than long-lived proteins or vice versa. In order to look for this specific enrichment, we broke the turnover ratio data into 5 bins ranging from very high turnover (quick half-life) to low turnover (long half-life). We then examined how many genes in each bin were differentially expressed (DEGs) in SCZ and normalized this to the number of genes in the bin, yielding the % of DEGs in each bin (Fig 2). A one-way ANOVA (F(4,10)=0.003; p>0.999) revealed that there was no particular enrichment in any one bin, suggesting that SCZ does not specifically enrich genes based on turnover ratio.

**Figure 2:**
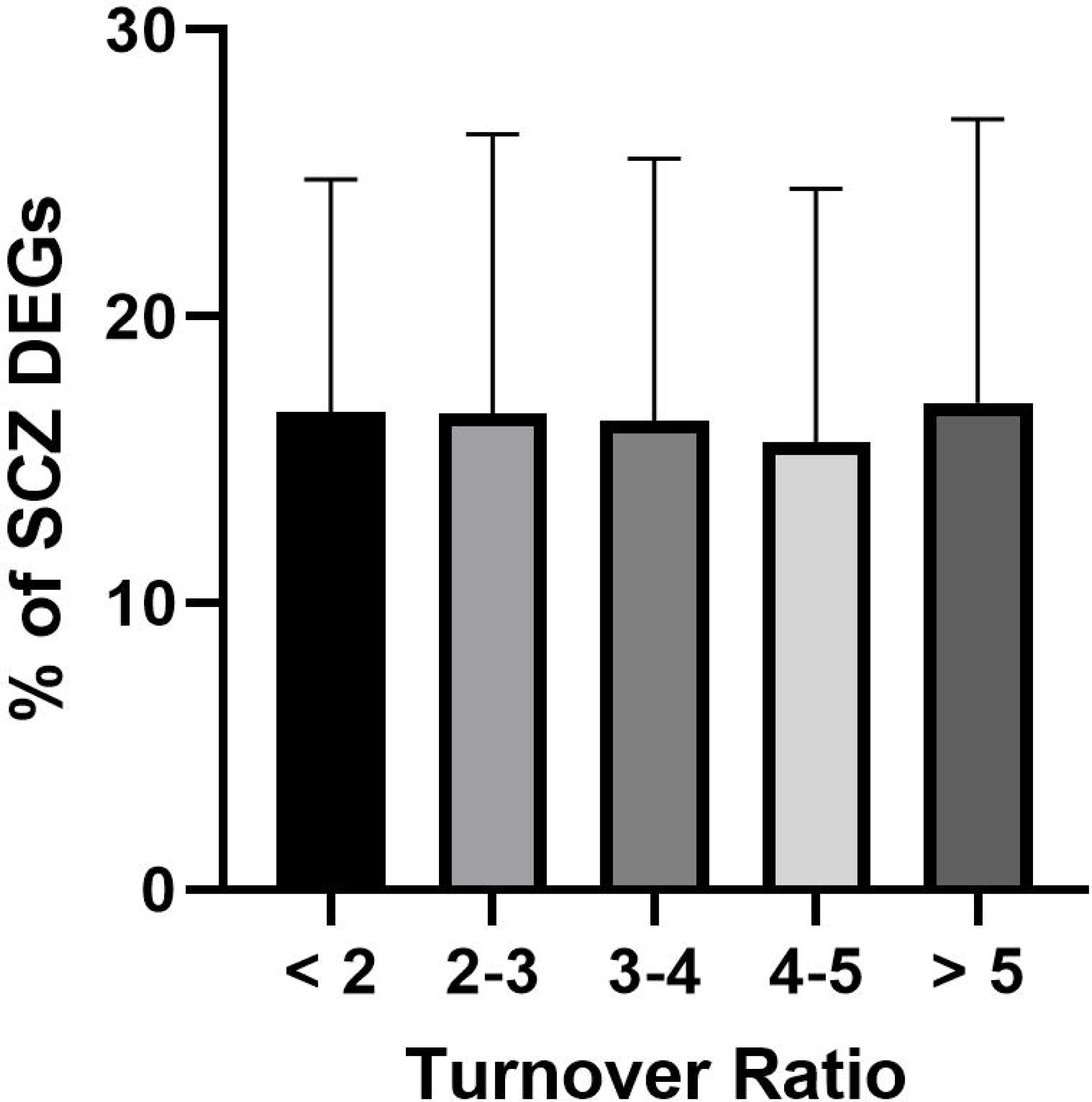
No shift in gene fold change in SCZ as a function of turnover ratio. Genes in different turnover ratio bins showed similar degrees of expression changes between SCZ and CTL samples across all 3 datasets, suggesting that no particular bin was especially sensitive to SCZ disease effects.

### High and low persistence genes correspond to pathways known to be impacted by SCZ

Turnover ratios were available for 2272 genes [13]. Of these, 2101 were found in our 3 RNA expression datasets, so persistence scores were calculated for these genes (Supplemental Table 1). The overall relationship between persistence and p value in our 3 SCZ datasets can be found in Fig 3. Genes that were significantly differentially expressed between SCZ and CTL and had a persistence score >0.5 or <-0.5 were determined to have high or low persistence, respectively. This 0.5 cutoff marked genes with good separation from the rest of the dataset, and therefore suggests that these genes have especially meaningful persistence. At this cutoff, we identified 30 high persistence genes (4 in Deep, 15 in DISC1, and 11 in Super) and 7 low persistence genes (0 in Deep, 5 in DISC1, and 2 in Super) (Table 1). We did not find much direct overlap between datasets (3 shared high persistence and 0 shared low persistence).

**Table 1:** “Persistent” genes in SCZ datasets. Genes that are significantly differentially expressed in SCZ vs CTL and have high (>0.5) or low (<-0.5) persistence scores.

| Deep |  |  |  | DISC1 |  |  |  | Super |  |  |  |
| --- | --- | --- | --- | --- | --- | --- | --- | --- | --- | --- | --- |
| High Persistence |  | Low Persistence |  | High Persistence |  | Low Persistence |  | High Persistence |  | Low Persistence |  |
| Gene | Score | Gene | Score | Gene | Score | Gene | Score | Gene | Score | Gene | Score |
| TNR | 0.663 |  |  | CRYM | 0.870 | ALDH1A1 | -0.567 | NEFM | 0.870 | UQCRB | -0.594 |
| ME3 | 0.625 |  |  | GPD1 | 0.791 | CLYBL | -0.594 | MBP | 0.791 | CORO2A | -0.631 |
| HIBADH | 0.600 |  |  | MBP | 0.750 | CHL1 | -0.631 | TUBB2A | 0.750 |  |  |
| CAPZA1 | 0.574 |  |  | TNR | 0.698 | NRN1 | -0.875 | NEFL | 0.698 |  |  |
|  |  |  |  | SYNPR | 0.670 | ATP1A4 | -0.960 | ATP1B1 | 0.631 |  |  |
|  |  |  |  | NCAN | 0.663 |  |  | INA | 0.625 |  |  |
|  |  |  |  | SLC32A1 | 0.631 |  |  | HSPD1 | 0.600 |  |  |
|  |  |  |  | KCNA1 | 0.625 |  |  | GNAO1 | 0.592 |  |  |
|  |  |  |  | SLC13A5 | 0.609 |  |  | SH3BGRL2 | 0.574 |  |  |
|  |  |  |  | NRGN | 0.600 |  |  | ATP6V0A1 | 0.561 |  |  |
|  |  |  |  | BDH1 | 0.592 |  |  | SEPT7 | 0.512 |  |  |
|  |  |  |  | ME3 | 0.574 |  |  |  |  |  |  |
|  |  |  |  | ACTN2 | 0.561 |  |  |  |  |  |  |
|  |  |  |  | PLCXD3 | 0.525 |  |  |  |  |  |  |
|  |  |  |  | GNG4 | 0.512 |  |  |  |  |  |  |

**Figure 3:**
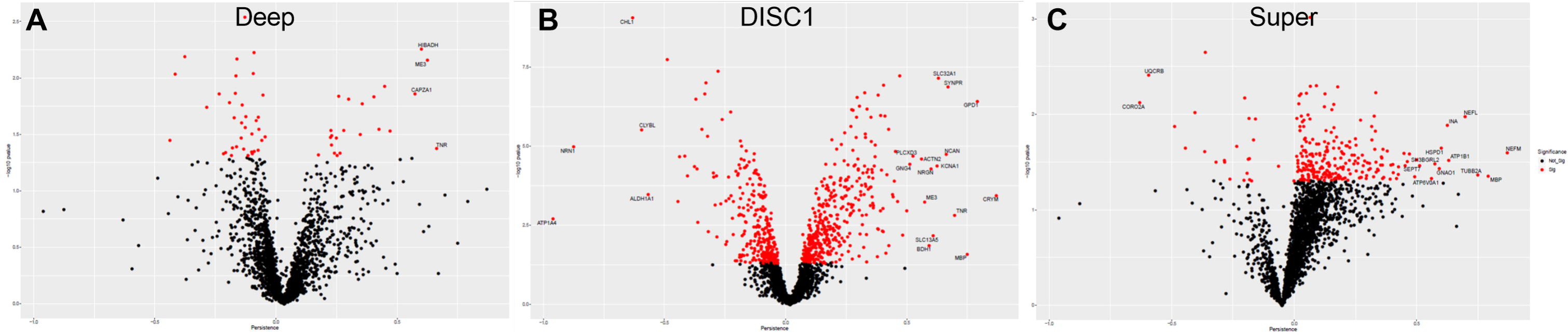
Persistent, significant genes in SCZ. Volcano plots of persistence scores and – log10 p values from (A) Deep, (B) DISC1, and (C) Super datasets. Red dots signify significant (p<0.05) genes, and labels signify genes that also surpassed the persistence cutoff of 0.5.These genes were assigned as having low (<-0.5) or high (>0.5) persistence and used in subsequent analyses.

We then used EnrichR to run a pathway analysis on the high and low persistence genes identified in each dataset [14]. Each set of genes was analyzed for significant (p < 0.05) enrichment in the gene ontology biological pathway, molecular function, and cellular component categories. Overall, we found 412 high persistence pathways (113 in Deep, 126 in DISC1, 149 in Super) and 59 low persistence pathways (0 in Deep, 39 in DISC1, 20 in Super). While these pathways were varied, there were a few themes that were common between datasets that are known to be impacted by SCZ [15,16]. The predominant theme in the high persistence genes was ion homeostasis, while the predominant low persistence theme was respiration. Other high persistence themes include metabolic process, immune system process, and transport; while other low persistence themes included ion homeostasis, ATP, and protein modification (Fig 4; Supplemental Table 2).

**Figure 4:**
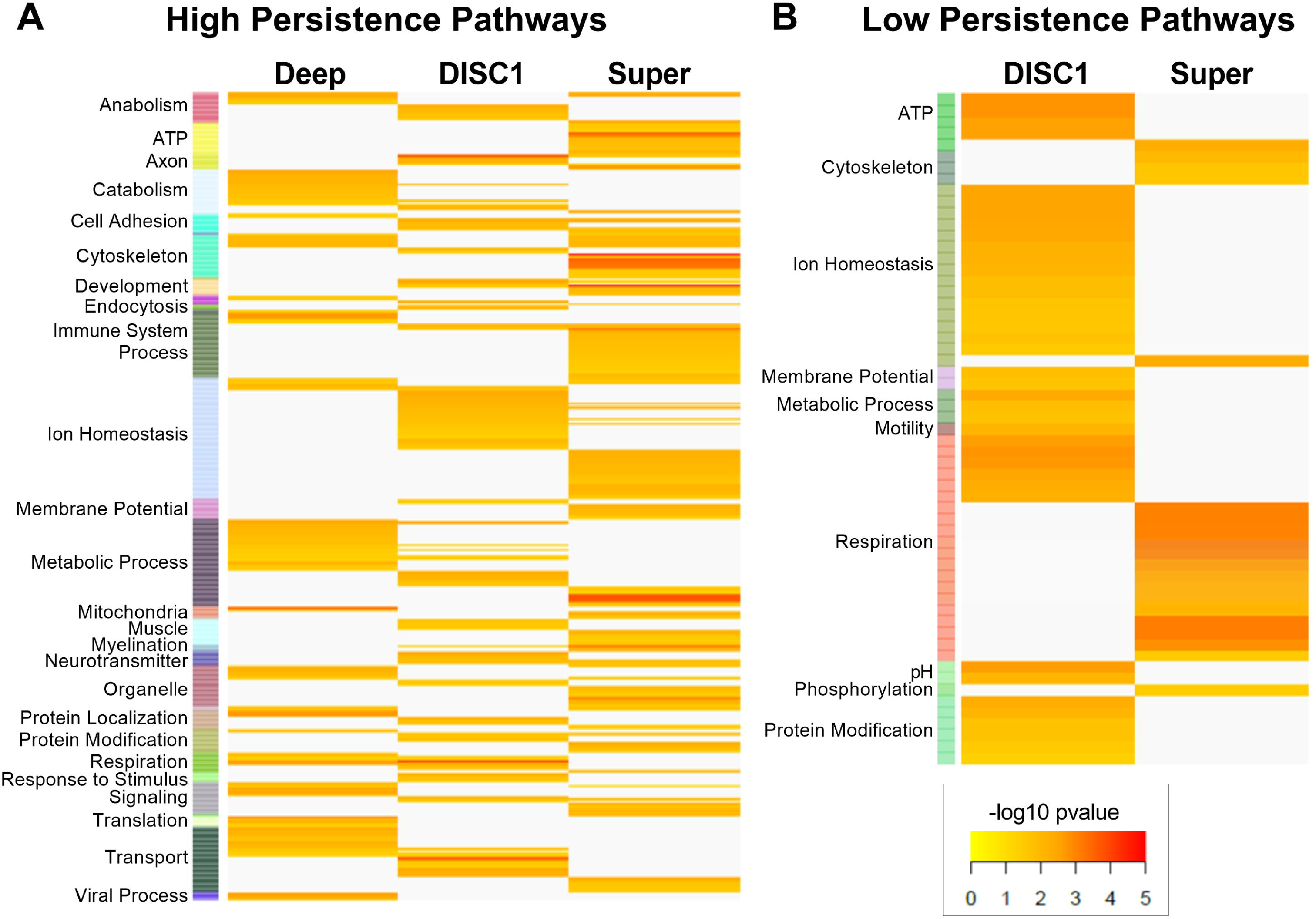
Persistent pathways in SCZ. Heatmaps of significant (p<0.05) pathways associated with (A) high and (B) low persistence genes. Pathways have been assigned to categories summarizing their major functions.

### Persistence improves RNA prediction of protein fold change

While the present persistence calculation appears to reveal meaningful insights into SCZ, we can compare our persistence scores to actual protein abundance measures to further demonstrate the value of this approach. The DISC1 dataset contains information on both RNA and protein fold changes, allowing for this insightful comparison. While persistence did not generate a perfect correlation with protein fold change (Pearson Correlation R^2^=0.65), it did generate a stronger correlation with protein fold change than mRNA fold change alone (Pearson Correlation R^2^=0.208) (Fig 5). This comparison suggests that persistence does improve our ability to predict protein abundance from RNA abundance. While there is still room for improvement, the persistence calculation presented here represents a good starting point for improving our ability to estimate changes in protein from changes in RNA.

**Figure 5:**
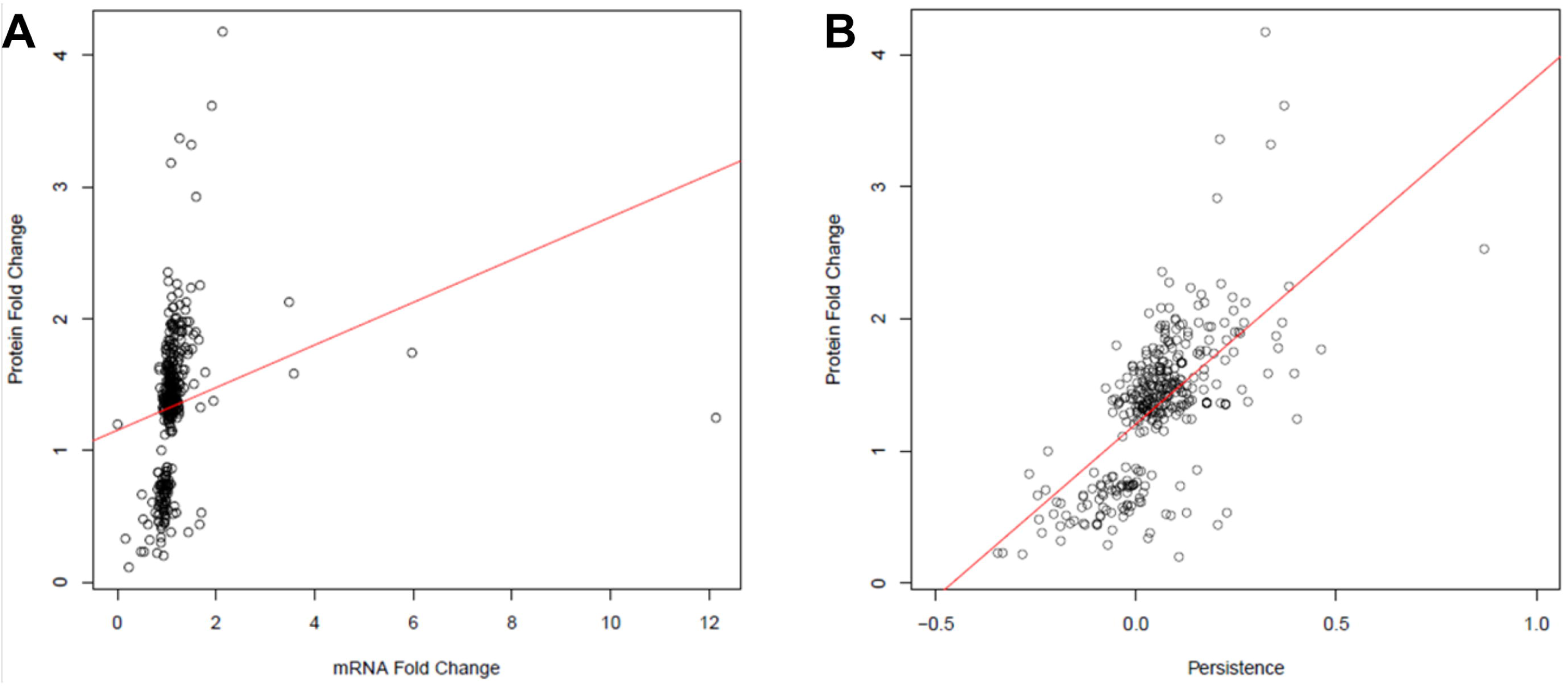
Persistence better predicts protein fold change in SCZ than mRNA fold change alone in the DISC1 dataset. (A) Correlation between protein and mRNA fold change (R^2^=0.208). (B) Correlation between protein fold change and persistence scores (R^2^=0.68). This improved correlation suggests that persistence can be predictive of protein expression but that room for improvement exists in the present calculation.

## DISCUSSION

Persistence is a novel concept intended to expand the lens of transcriptomics to infer protein function by considering RNA expression and protein turnover together. Our goal is to address a common problems with transcriptomics, in that it tries to gain functional insight into protein expression that is often missed by simply analyzing RNA expression [6–8]. We propose that one can approximate protein abundance by considering RNA expression in conjunction with how long the corresponding protein will last in the cell once it is translated (turnover ratio) [11,12]. While there is still room for improvement to make this link more complete between RNA and protein, we have shown that persistence is an insightful measure and that further work with the concept could be a valuable resource for computational biologists.

The lack of relationship between protein turnover and RNA abundance suggests that a more complex metric is required to extract meaningful insights from the data (Fig 1). For this reason, we developed the concept of persistence and demonstrated how it may be applied in 3 SCZ RNA datasets (1 RNAseq and 2 microarray). We did not observe a particular enrichment of high or low turnover genes in SCZ relative to CTL (Fig 2). This suggests that there is not one specific type of turnover impacted by SCZ, but does not rule out that enrichment of certain high or low persistence genes happens in the disease.

Therefore, we identified high and low persistence genes by selecting significantly differentially expressed genes with a persistence score >0.5 or <-0.5, respectively (Fig 3; Table 1). These criteria yielded a small number of genes that largely corresponded to pathways known to be altered in SCZ (Fig 4). These results are encouraging that persistence is a meaningful measure because these pathways have been implicated in SCZ in prior studies. For example, ion homeostasis was the most commonly altered category in our high persistent pathways [17–22]. Specifically, potassium and sodium metal ion activity and transport appear to be more persistent in SCZ. Multiple studies in animals, iPSCs, and humans have noted increased potassium and sodium channel expression and activity in the prefrontal cortex. These changes were associated with abnormal neuronal activity, diminished synaptic plasticity, and impaired white matter integrity, all of which are characteristic of SCZ [17–22]. Additionally, antipsychotics can reverse these ion channel alterations, suggesting that this is a key mechanism in SCZ pathology [23–25]. Beyond ions, we also saw increased persistence in inflammatory system processes, which is consistent with observations of increased cytokine expression and immune system responsivity in SCZ [20,26–28].

In terms of low persistence, we observed a decrease in respiration in SCZ. Specifically, there was low persistence in pathways associated with oxidative phosphorylation, suggesting that neurons in SCZ struggle to maintain sufficient levels of ATP production via aerobic respiration. Impaired oxidative phosphorylation and abnormal mitochondrial function has been noted in multiple SCZ studies [29–32]. This again indicates that there is a loss of efficiency in SCZ that puts more stress on the system and forces neurons to turn to alternate sources of energy. Indeed, postmortem samples from SCZ patients show an increase in lactate metabolism [29,33]. Interestingly, we also observed high persistence in metabolic processes, which further supports this concept of metabolic compensation for a loss of oxidative phosphorylation in SCZ. Overall, the genes and pathways identified by our persistence analysis are altered in SCZ, supporting the potential of this technique to extract important functional information from transcriptomics datasets.

### Limitations and Future Directions

The present persistence results did correlate fairly strongly with protein abundance (R^2^=0.68), which was a substantial improvement over RNA expression alone (R^2^=0.208). However, continued efforts to improve the persistence calculation are warranted (Fig 5). We were only able to do this comparison in one dataset, as matching postmortem or iPSC tissue RNAseq and proteomics is rare, but this suggests that there is room for improvement in our persistence calculation. The present study was also limited by the narrow scope of the available turnover ratios. We only had ratios for ∼2200 proteins, and these ratios were specific to synaptosomes. This left us with a narrow window into RNA datasets that needed to be specifically from neurons [34]. Of our more than 30 SCZ datasets, only 3 matched these criteria. With limited RNAseq / microarray datasets and turnover ratios, the present calculation is a good concept demonstration, but we would like to develop the concept of persistence further using expanded datasets and machine learning.

Ideally, this expansion would feature well-powered studies that have RNAseq and proteomics run on the same samples. This would allow us to use machine learning to integrate RNA expression and protein turnover ratios in the way that would most accurately predict protein expression in the same tissue [35,36]. A more complex relationship undoubtedly exists between these factors, so training a model to more accurately predict this relationship would be highly beneficial [10–12]. It would also be useful to expand the number and variety of protein turnover ratios to put into this model. This would require more SILAM studies [37,38], but gathering this information from more animals in more tissues would greatly expand the context in which persistence could be applied. For example, another tissue for which turnover data is currently available is the liver [13]. We could use RNAseq data and liver cell protein turnover ratios to create models to predict protein abundance in a normally functioning liver. This model could then be used with RNAseq data from liver cancer to better identify the functional changes that occur in that disease state. While this would require time and money, it would add a new dimension to transcriptomics studies and improve our ability to understand mechanisms underlying various disorders. Especially high or low persistence genes may also represent important therapeutic targets since they will likely have a magnified role in disease mechanisms [39–41].

Overall, we have demonstrated that applying protein turnover ratio data to RNA expression data represents a novel form of analysis that expands the amount of information that can be obtained from transcriptomics. Persistence combines RNAseq with protein turnover ratios to infer protein abundance, which is a more accurate measure of function in a cell. While the present study is limited by a lack of sufficient input data, it does identify high and low persistent genes and pathways that have been implicated in SCZ. Further development of the concept of persistence with expanded studies and machine learning techniques could greatly improve our ability to understanding of the molecular landscape in disease with RNAseq alone.

## MATERIALS AND METHODS

### Dataset Selection

Given that the turnover ratios were derived from synaptosomes, the present analysis was restricted to RNA datasets from neuronal populations alone. While this concept could be applied to a variety of subjects, we selected SCZ as our present focus, as it is a particularly pervasive disorder known to have widespread effects throughout the brain. With these criteria in mind, we selected three neuronal, SCZ RNA datasets: Deep, DISC1, and Super [34]. The Deep and Super datasets were derived from pyramidal neurons cut from the dorsolateral prefrontal cortex of postmortem SCZ and CTL samples using laser capture microscopy. Specifically, the Deep dataset comes from the deep layers (IV-VI) of this region while the Super dataset comes from the superficial layers (I-III). Both of these datasets were obtained from RNA microarrays. The DISC1 dataset was derived from RNAseq run on induced pluripotent stem cells (iPSCs) obtained from SCZ patients positive for a DISC1 mutation and CTL siblings lacking the mutation. These iPSCs were differentiated into neurons and used for RNAseq analysis. Additionally, the DISC1 dataset included parallel proteomic data (mass spectroscopy) which allowed us to compare mRNA and protein.

### Turnover vs RNA Abundance

Scatterplots of protein turnover versus mRNA abundance we generated utilizing R base graphics for the Deep, DISC1, and Super datasets. This was performed for both CTL and SCZ data. A linear regression model was computed for both groups of data using the R base statistics *lm* function.

### Turnover Distribution in SCZ

Genes were sorted into bins of low to high turnover ratios. Differentially expressed genes (DEGs) (p<0.05) in SCZ were overlaid with these bins to determine the number of genes in each turnover bin. One-way ANOVA was used to determine if there was any significant difference between the numbers of DEGs in each bin, which would be suggestive of a particular bin being more strongly affected by SCZ. This analysis was performed in GraphPad Prism 8.

### Persistence Calculation

In order to identify genes with particular importance, we created the concept of persistence. Persistence utilizes the Huganir turnover ratios [13] and RNA fold change to generate a measurement of the potential functional impact of changes in RNA expression. The persistence calculation was conducted as follows:

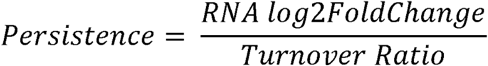

This equation is designed to generate a persistence score in which high values indicate high persistence, meaning that that particular gene is produced more in the disease state (high RNA fold change) and stays around longer to exert more activity (low turnover ratio [i.e. long half-life]). On the other hand, low persistence suggests that genes are produced less (low RNA fold change) and are quickly degraded (high turnover ratio [i.e. short half-life]), resulting in a lower overall impact in the synaptosome. Theoretically, this combination of RNA fold change and turnover ratio should act as a proxy for protein fold change.

### Volcano Plots

The persistence scores for the Deep, DISC1, and Super datasets were generated and quantile normalized using the *preprocessCore* library in R. The negative log 10 of the p-value was computed for SCZ versus CTL. A volcano plot was then generated using the *ggplot2* R package with persistence on the x-axis and log10 p-value on the y-axis. Significant genes were colored in red with the threshold of 0.05 for the p-value or 1.3 for -log10 of the p-value. Those genes which were significant were further filtered for persistence. The thresholds for persistence were greater than 0.5 or less than -0.5. The genes that were selected were then used for downstream analysis.

### Pathway Enrichment Analysis

After selecting significant, persistence genes, these genes were then used for enrichment analysis utilizing EnrichR [14]. The databases utilized were the three Gene Ontology databases: Cellular Component, Biological Process, and Molecular Function. The analysis was performed for the set of genes with high and low persistence separately, yielding a set of enriched pathways. A heatmap of -log10 p-values was generated using the library *gplots* for the Deep, DISC1, and Super datasets utilizing the union of significant pathways for the three datasets. The resulting pathway annotations were then categorized into supersets using *a priori* knowledge.

### DISC1 mRNA vs Protein Fold Change

R base graphics were used to generate a mRNA log2 fold change versus protein log2 fold change scatterplot for the DISC1 dataset. Correlation was computed using R base statistics and a linear regression model was fitted using R base statistics.

### DISC1 Persistence vs Protein Fold Change

R base graphics were used to generate a QN persistence versus protein log2 fold change scatterplot. As in previous method descriptions, correlation was computed using R base statistics and a linear regression model was fitted.

## AUTHOR CONTRIBUTIONS

MAS, JKR, and REM developed the concept. MAS and JKR conducted the analyses and wrote the manuscript. REM reviewed and edited the manuscript.

## SUPPORTING INFORMATION LEGENDS

**Supplemental Table 1: Persistence information for full SCZ datasets**. Persistence scores, -log 10 p values, and significance information for 2101 genes in Deep, DISC1, and Super dataset.

**Supplemental Table 2: Full pathway information from Figure 4 heatmaps**. Details regarding pathways associated with (A) high and (B) low persistence genes.

